# A vacuolar invertase gene *SlVI* modulates sugar metabolism and postharvest fruit quality and resistance in tomato

**DOI:** 10.1101/2024.06.20.599942

**Authors:** Yu Wu, Haonan Chen, Mengbo Wu, Yuanyi Zhou, Chuying Yu, Qihong Yang, Filip Rolland, Bram Van de Poel, Mondher Bouzayen, Nan Hu, Yikui Wang, Mingchun Liu

## Abstract

Sugars act as signaling molecules to modulate various growth processes and enhance plant tolerance to various abiotic and biotic stresses. Moreover, sugars contribute to the post-harvest flavor in fleshy fruit crops. To date, the regulation of sugar metabolism and its effect in plant growth, fruit ripening, postharvest quality and resistance remains not fully understood. In this study, we investigated the role of tomato gene encoding a vacuolar invertase, hydrolyzing sucrose to glucose and fructose. *SlVI* is specifically expressed during the tomato fruit ripening process. We found that overexpression of *SlVI* resulted in increased leaf size and early flowering, while knockout of *SlVI* led to increased fruit sucrose content, enhanced fruit firmness, elevated resistance of postharvest fruit to *Botrytis cinerea.* Moreover, the content of naringenin and total soluble solids was significantly increased in *SlVI* knockout fruit at postharvest stage. Transcriptome analysis showed a negative feedback regulation triggered by sucrose accumulation in *SlVI* knockout fruit resulting in a downregulation of *BAM3* and *AMY2,* which are critical for starch degradation. Moreover, genes associated with cell wall, cutin, wax, and flavonoid biosynthesis and pathogen resistance were upregulated in *SlVI* knockout fruit. Conversely, the expression levels of genes involved in cell wall degradation were decreased in knockout fruit. These results are consistent with the enhanced postharvest quality and resistance. Our findings not only provide new insights into the relationship between tomato fruit sucrose content and postharvest fruit quality, but also suggest new strategies to enhance fruit quality and extend postharvest shelf life.

## Introduction

Sucrose serves as a direct source of carbon and energy for plant metabolism and storage, but it can also act as a signaling molecule that interacts with different phytohormone signaling networks (Petreikov et al., 2009; Eveland and Jackson, 2011). Through these interactions, sucrose plays a pivotal role in regulating various cellular processes crucial for plant growth and development. These include, but are not limited to, embryo establishment, seed germination, lateral root induction, and meristem development, notably influencing cell division and expansion (Riou et al., 2000; Tognetti et al., 2013) Moreover, sugars play a central role in stress perception and signaling, modulating stress-induced gene expression and directing carbon allocation to adjust osmotic balance and scavenge reactive oxygen species. These integrated processes collectively aid in alleviating the detrimental effects of environmental stress on plant physiology (Saddhe et al., 2021; Gautam et al., 2022; Jeandet et al., 2022).

Sugar metabolism in plants is a dynamic and complex process involving the loading, unloading, and long-distance transport (source to sink) of sugars. This process is largely controlled by sugar transporters and hydrolases (Remi et al., 2013).The translocation of sugars from source tissues (e.g., leaves) to sink tissues (e.g., fruit) primarily involves passive mass flow within the phloem. However, active mechanisms are necessary for loading and unloading, requiring the transport of sugars across cell membranes. This transport is facilitated by various types of transporter proteins, including SUC/SUT sucrose-H+ symporters, proton-coupled MST (monosaccharide sugar transporter), and SWEET (Sugars Will Eventually be Exported Transporter) uniporters (Braun et al., 2014; Shammai et al., 2018; Sun et al., 2022a). In sink tissue, sucrose is hydrolyzed into hexoses (glucose and fructose) or their nucleotide derivatives (UDP/ADP-glucose and fructose) through the enzymatic actions of invertases (INV) and sucrose synthases (SUS/SuSy) respectively, facilitating participation in diverse metabolic processes (Sturm, 1999; Ruan et al., 2010). Invertases can be classified into different isoforms, including acidic vacuolar invertases (VIN), cell wall invertases (CWIN), and neutral cytoplasmic invertases (CIN), based on their optimal pH, solubility, and subcellular localizations. Imported or symplastically unloaded sucrose can be hydrolyzed by CIN in the recipient cytoplasm or transported into vacuoles for subsequent hydrolysis into glucose and fructose by VIN (Lu et al., 2009; Sun et al., 2022b). Intracellular glucose and fructose can be allocated to various metabolic processes such as glycolysis and respiration, the synthesis of polymers such as cellulose and starch, or storage in vacuoles. They can also act as signaling molecules, thereby modulating gene expression (Rolland et al., 2006; Lastdrager et al., 2014). The activities of VIN and CWIN are also modulated post-translationally by inhibitor proteins (Jin et al., 2009; Zhang et al., 2015; Qin et al., 2016).The tomato fruit is an important sink organ with a very dynamic sugar metabolism. During the early stages of tomato fruit development, for example, there is an important transient accumulation of starch that can reach levels of up to 20% of the fruit’s dry weight (Nguyen-Quoc and Foyer, 2001). During fruit ripening, starch stored in the plastids is broken down into sugars, contributing to the soluble hexose level in the mature fruit (Carrari and Fernie, 2006). This transition, known as the starch-sugar interconversion, is facilitated by enzymes such as α-amylases and invertases (Nicolas et al., 2023; Yin et al., 2023). Glucose, fructose, and sucrose are the primary sugars found in ripe tomato fruit. Consequently, the accumulation of soluble sugars and their relative ratios play a significant role in shaping the flavor of postharvest tomatoes (Kader, 2008; Oms-Oliu et al., 2011).

In recent years, studies employing RNA interference (RNAi) have provided valuable insights into the specific genes involved in sucrose metabolism and transport and their impact on various aspects of tomato plant physiology. Suppression of *SUS1* in tomato resulted in a noticeable reduced growth rate and reduced fruit set shortly after flowering. Besides, *SlSUS*-RNAi lines showed altered expression of genes associated with leaf morphology and auxin levels (Goren et al., 2017). Similarly, the targeted suppression of the cell wall invertase *LIN5*, led to reduced glucose and fructose levels and detrimental effects on reproduction, including aberrant pollen morphology, decreased pollen viability and germination rate, and a decline in fruit set, size, and seed number (Zanor et al., 2009). This highlights the importance of sucrose hydrolysis in normal fruit development and fertility. In addition, functional studies on sugar transporters have also shown that inhibition of *SlSUT1* led to sucrose and starch accumulation in leaves due to blocked phloem loading, triggering premature leaf senescence while suppression of *SlSUT2* only affected tomato fruit and seed development with deficient pollen tube elongation, leading to the production of small, sterile fruit with a reduced seed number. Additionally, silencing of *SlSUT4* increased leaf sucrose export and increased drought tolerance of tomato plants (Hackel et al., 2006; Liang et al., 2023).RNAi-mediated suppression of hexose transporters, such as *LeHT3*, led to a significant reduction in hexose (glucose and fructose) concentrations within the fruit, while photosynthetic rates and sucrose unloading from the phloem in leaves remained unaffected (Gear et al., 2000; Mccurdy et al., 2010). Surprisingly, RNAi silencing of *SlSWEET7a* and *SlSWEET14*, which encode plasma membrane localized transporters, resulted in taller plants and larger fruit with enhanced sugar accumulation in mature fruit. This effect could be attributed to the observed increase in invertase activity and expression of other SWEET members (Zhang et al., 2021). Overexpression of *SlSWEET11b* led to the redistribution of sugars in the stem, increases stem diameter, and accumulates more lignin. Further studies showed that the size, weight and sugar content of fruit were significantly increased (Sun et al., 2023). However, the specific role of the vacuolar invertases SlVI in regulating sucrose content and its impact on tomato plant physiology and postharvest quality of fruit are still not fully understood.

In this study, through integration of two transcriptome datasets, we revealed that the *SlVI* gene displays a distinct expression pattern during the tomato fruit ripening process. To further elucidate the role of SlVI in tomato plant growth and fruit quality, we conducted experiments involving overexpression of the *SlVI* gene and employed CRISPR/Cas9-mediated gene editing to generate *SlVI* knockout lines. Our analyses showed that overexpression of the *SlVI* not only resulted in increased chlorophyll content and leaf size, but also accelerated flowering and reproductive growth. Conversely, *SlVI* knockout lines produced fruit with higher soluble solids and naringenin chalcone content. Furthermore, *SlVI* knockout fruit exhibited higher firmness and extended shelf life. Gray mold infection demonstrated that *SlVI* knockout fruit displayed enhanced resistance to *Botrytis cinerea* compared to the wild-type. Moreover, the content of cellulose, hemicellulose, and protopectin (three main components of the cell wall) significantly increased in the *SlVI* knockout fruit. Further investigation revealed that the *SlVI* knockout pericarp exhibited a smaller and denser cell structure based on staining of paraffin transverse sections of tomato fruit. These findings collectively contribute to our understanding of the link between tomato fruit sucrose content and postharvest fruit quality.

## Results

### Tomato vacuolar invertase gene *SlVI* is specifically expressed during fruit ripening process

Starch serves as the principal storage carbohydrate in plants and plays a crucial role in modulating sugar homeostasis, particularly in response to fluctuating environments. Some fruits store starch early in fruit growth and undergo degradation during the ripening progress (MacNeill et al., 2017; Roch et al., 2019). To investigate the dynamics of starch and sugar content during fruit development and ripening, we utilized liquid chromatography and mass spectrometry (LC-MS) to measure these compounds at various developmental stages, ranging from 10 days post-anthesis (DPA) to 30 DPA, as well as different ripening stages: breaker (Br), Br + 3 (3 days post-breaker), Br + 5, Br + 7, and Br + 10 in Micro Tom tomato (Fig. 1A and S1). The results reveal two distinct phases of starch accumulation in tomato fruit. In the first phase (10-20 DPA), corresponding to fruit development and rapid growth, starch gradually accumulates, reaching its peak at 20 DPA, and then gradually degrades. During the second phase (27-30 DPA), another peak of starch accumulation is observed at 30 DPA, followed by degradation at the onset of ripening (Fig. 1A). Interestingly, the three main sugars also exhibit two accumulated peaks, occurring at 25 DPA and Br+5 which coincide with the stages of starch degradation and consistently lag behind the peak of starch accumulation (Fig. 1B). These results suggest that tomato fruit initially stores carbon as starch during the early rapid growth stage and subsequently mobilizes it during later development and ripening stages. Additionally, the simultaneous occurrence of starch degradation and sugar accumulation underscores the dynamic balance and transformation process between starch and sugar during tomato fruit development and ripening.

**Figure 1.**
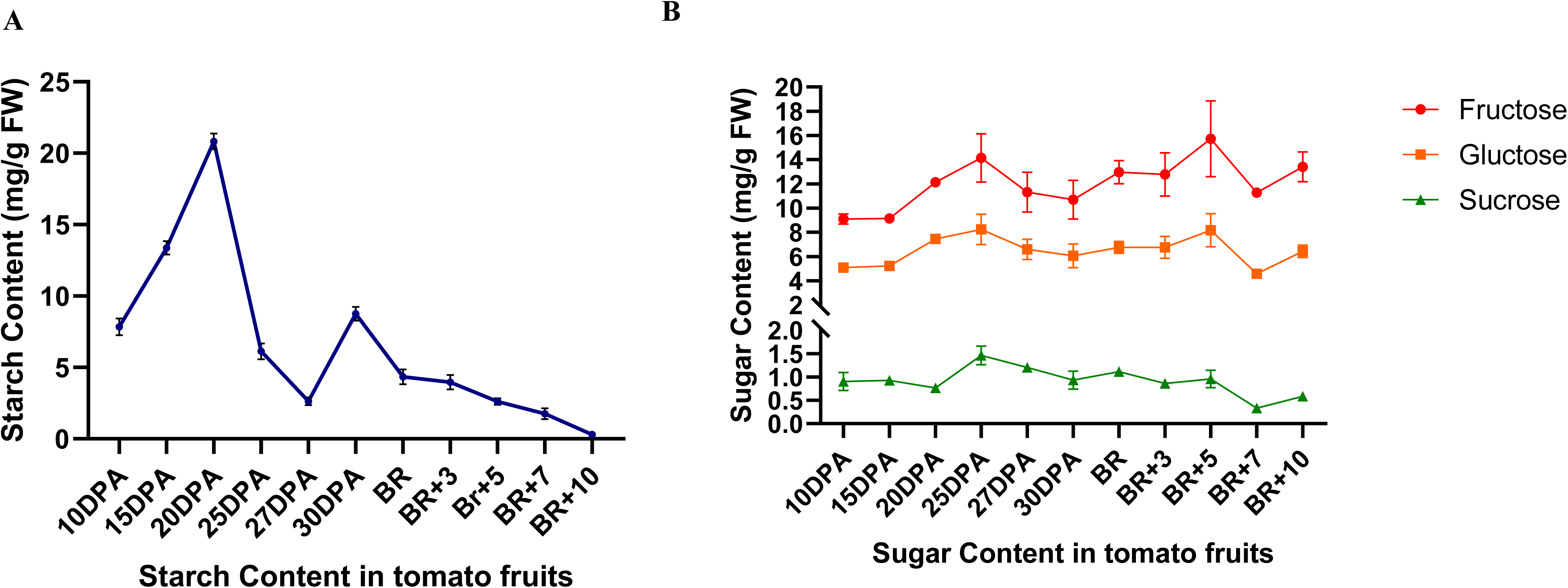
Starch degradation and sugar content changes during Micro Tom tomato fruit development. A) Starch content at different fruit developmental and ripening stages of tomato fruit B) Sugar content at different fruit developmental and ripening stages of tomato fruit. Under the same growth conditions, six fruit were collected in each stage, mixed and ground as samples for determination, and each of them has three technical repetitions.

To further explore the metabolism and transport of sugar during tomato fruit ripening, we conducted an analysis of the expression of genes related to sucrose transport and metabolism in the transcriptome datasets of Micro Tom and revealed a specific increase in expression of a vacuolar invertase gene (*SlVI*, *Solyc03g083910*) during fruit ripening (Fig. 2A, S2, S3) (Li et al., 2020). Phylogenetic analysis showed the presence of just two vacuolar invertases, SlVIN1/VI and SlVIN2/LIN9 in tomato (Fig. S4). To validate the specific expression pattern of *SlVI* during fruit ripening, RT-qPCR analysis was performed in various tissues including root, stem, leaf, bud, flower, and fruit at different developmental and ripening stages. The results of the RT-qPCR analysis showed that *SlVI* displays high expression levels during fruit ripening stages (Fig. 2B). The combined evidence from the transcriptome data and tissue-specific expression analysis highlights the pronounced expression of *SlVI* during fruit ripening compared to other genes involved in sugar hydrolysis and transport. This observation suggests that *SlVI* may have an important role in regulating fruit sugar content.

**Figure 2.**
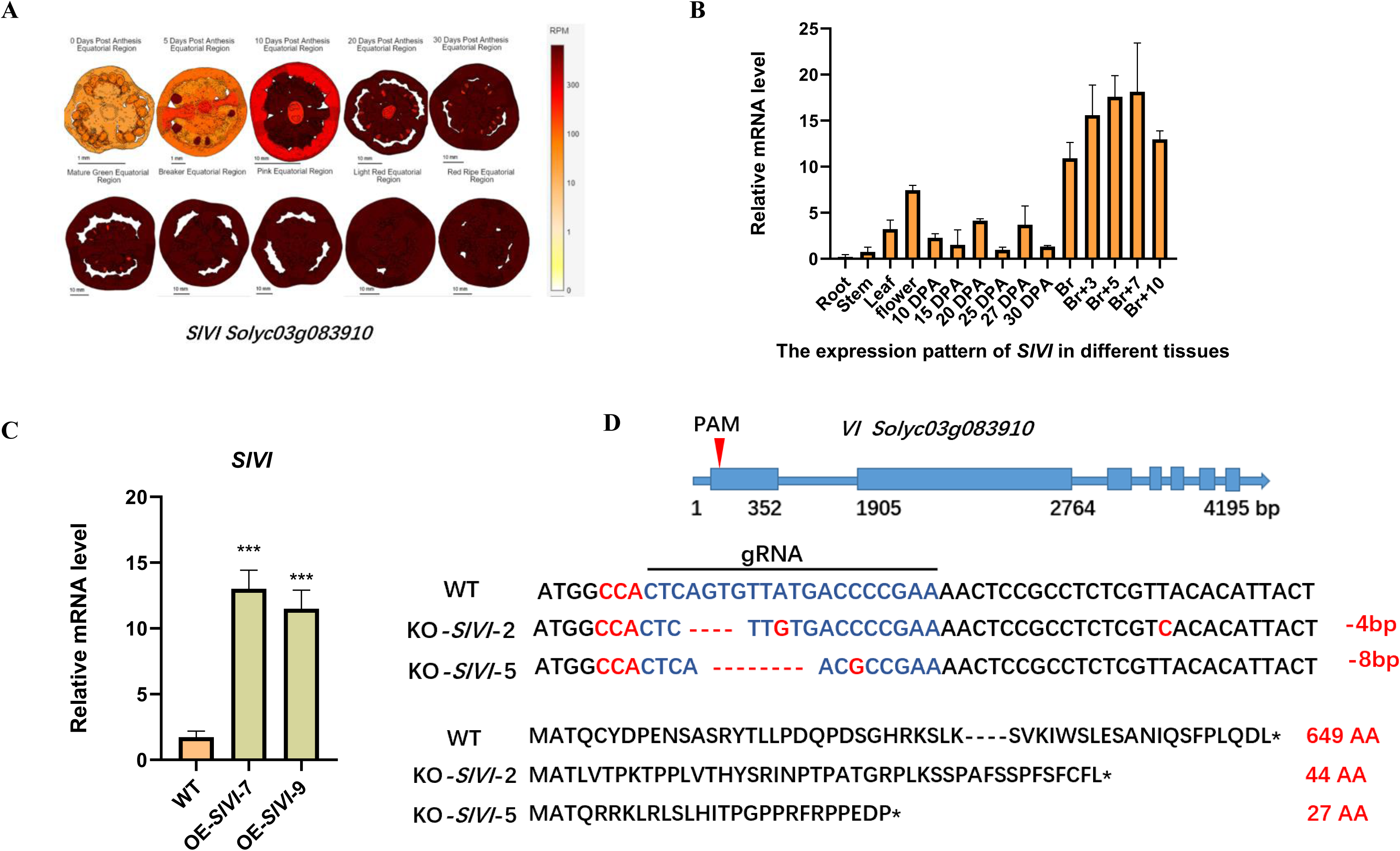
Tomato vacuolar invertase (*SlVI*) is specifically expressed during fruit ripening. A) Expression pattern images of *SlVI* gene from TEA *S.lycopersicum* M82 RNA-seq data. B) Relative *SlVI* transcript levels in different tissues and fruit development stages in Micro Tom wild-type tomato. DPA, days post anthesis; Br, fruit at breaker stage. C) Relative *SlVI* transcript levels in WT and T1 generation *SlVI* overexpression lines in tomato leaves. Data are presented as means ± SD (n = 3). Asterisks indicate statistical significance using Student’s test *p < 0.05, **p < 0.01 and ***p < 0.001 and ns indicates no significant change when compared with wild type. D) Schematic representation of CRISPR/Cas9-mediated gene editing at the first exon of the *SlVI* gene. Sanger sequencing showed two different base deletion mutations in T1 homozygous lines. The corresponding amino acid sequence indicated the production of a premature stop-codon and formation of truncated protein variants after editing. The asterisk represents the translation stop, the blue font indicates the sequence of the gRNA, the red font the PAM, deletions and mutations.

**Figure 3.**
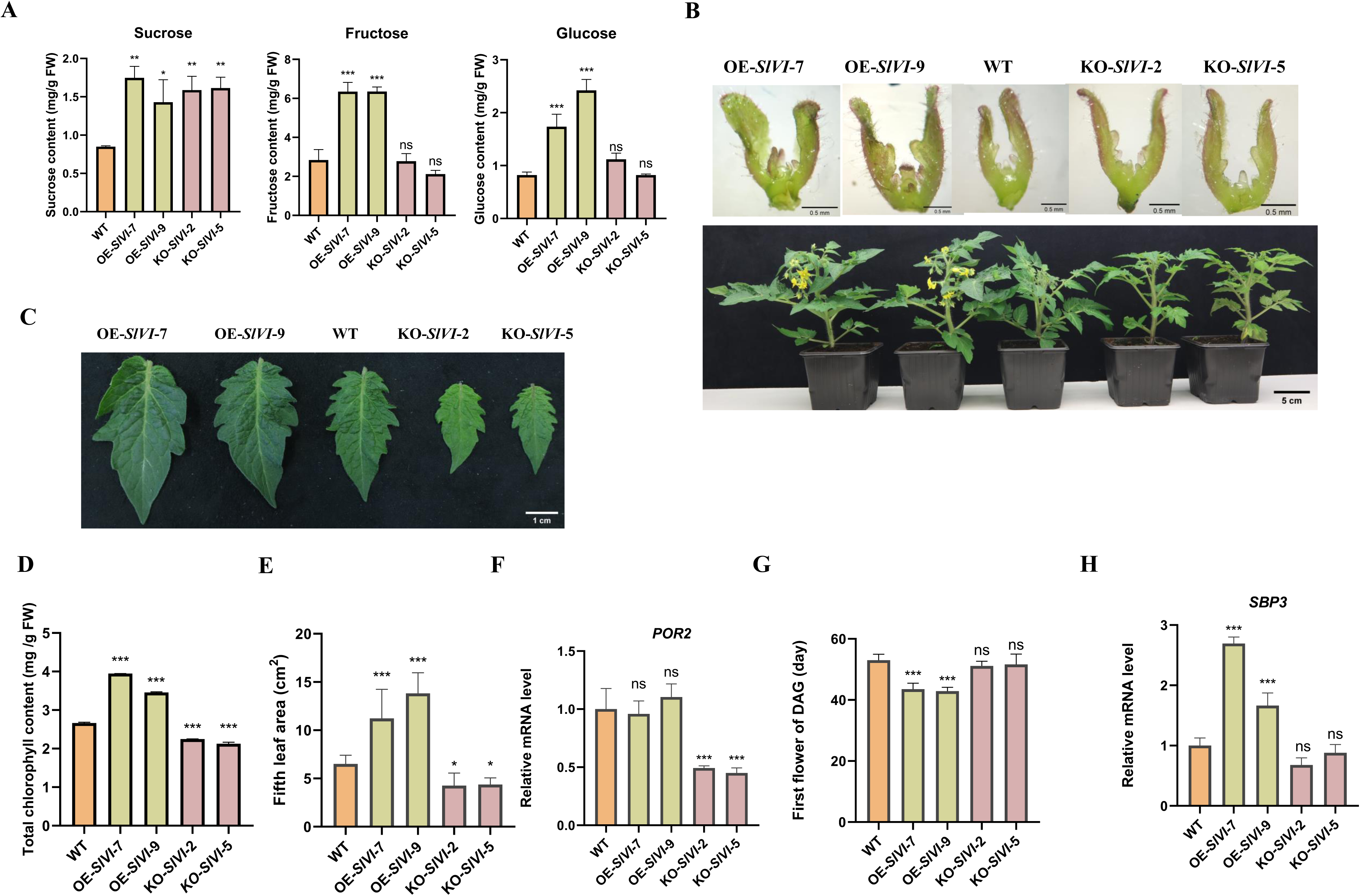
Overexpression of *SlVI* leads to earlier flowering and increased leaf size and chlorophyll content. A) Soluble sugar (sucrose, fructose, glucose) contents in WT, KO-*SlVI* and OE-*SlVI* leaves. B) Microscopic photos of flower primordia in seedlings ten days after germination (Bar = 0.5 mm) and the premature flowering phenotype of OE-*SlVI* plants (Bar = 5 cm). C) Areas of fifth leaf six weeks after germination. Leaves were photographed and areas measured by ImageJ. D) Total chlorophyll content in WT, KO-*SlVI* and OE-*SlVI* leaves. Asterisks indicate statistical significance using Student’s test *p < 0.05, **p < 0.01 and ***p < 0.001 and ns indicates no significant change, n=3. E) Phenotype of WT, KO-*SlVI* and OE-*SlVI* lines fifth leaves six weeks after germination. Bar = 1 cm. F) Relative *POR2* transcript levels in WT, KO-*SlVI* and OE-*SlVI* leaves. G) Time from germination to first flower in WT, KO-*SlVI* and OE-*SlVI* lines. H) Relative *SBP3* transcript levels in WT, KO-*SlVI* and OE-*SlVI* leaves. Data are presented as means ±SD. (a, n = 3 ;(c, n=20; (d, n=3;(e, n=3; (g, n= 20; (h, n=3). Asterisks indicate statistical significance using Student’s test *p < 0.05, **p < 0.01 and ***p < 0.001 and ns indicates no significant change when compared with wild type.

### Overexpression of *SlVI* leads to earlier flowering and increase leaf size and chlorophyll content

To elucidate the function of SlVI in tomato sugar metabolism and investigate its impact on plant physiology and development, we generated both overexpression and knockout lines. Two independent overexpression lines driven by a CaMV35S promoter (OE-*SlVI*-7, and 9) show a significant increase in mRNA level of *SlVI* in leaves compared to the wild type (WT) (Fig. 2C). Additionally, we obtained two Cas9-free homozygous knockout lines (KO-*SlVI*-2 and 5) by CRISPR/Cas9-mediated gene editing using a specific guide RNA targeting the first exon (Fig. 2D). Based on the Sanger sequencing results, it was shown that there were 4 and 8 base pair deletions in the two knockout lines, respectively. These deletions resulted in premature stop codons and truncated protein versions (Fig. 2D).

We first analyzed the effects of *SlVI* knockout and overexpression on the plant phenotype and sugar metabolism during vegetative growth. Surprisingly, both knockout or overexpression of this gene resulted in increased sucrose contents in tomato leaves. However, only the overexpression of *SlVI* significantly elevated the levels of sucrose hydrolysis products fructose and glucose (Fig. 3A). Interestingly, the overexpression lines exhibited darker green leaves and larger leaf size, while the knockout lines showed the opposite phenotype when compared with WT plants (Fig. 3C). Further analysis revealed that the chlorophyll content of overexpression leaves was significantly higher compared to WT. Conversely, the chlorophyll content in the knockout line was significantly lower than in WT leaves (Fig. 3D). Additionally, six weeks after germination, the fifth leaf of the tomato plant was selected for leaf size measurements. The results showed that compared to the WT, the leaves of the knockout plants were indeed significantly smaller, whereas the leaves of the overexpression plants were significantly larger (Fig. 3E). Consistently, expression of *POR2*, encoding protochlorophyllide oxidoreductase, a critical enzyme in chlorophyll synthesis, was downregulated in knockout lines, while no significant change was observed in overexpression lines (Fig. 3F). In addition to the above-mentioned phenotypes, we also observed that the flowering time of overexpression plants was earlier than that of wild-type plants. Specifically, the expression level of *SBP3*(encoding an SPL/SBP-box protein), which is essential for the plant to initiate flowering and control flowering time, was significantly upregulated in overexpression lines (Fig. 3B, G, H) (Silva et al., 2018). These findings suggest that altering sugar metabolism, specifically through the overexpression of *SlVI*, may enhance chlorophyll content and leaf size, and also shorten the transition from the vegetative to the reproductive phase in tomato.

### Knockout of *SlVI* significantly increases total soluble solids, sucrose and naringenin content in postharvest fruit

In addition to investigating the impact of *SlVI* expression on vegetative growth, we also examined its effects on fruit development and ripening. Both pictures and quantification of the fruit ripening process duration (the time interval from anthesis to the breaker stage) show that there is no significant difference between overexpression, knockout and wild type plants (Fig. 4A and S5). To understand the effect of altered *SlVI* gene expression on fruit flavor and nutritional quality, we analyzed changes in total soluble solids, sugar, organic acid, and flavonoid content in transgenic fruit. The total soluble solids content in *SlVI* knockout fruit was higher than in WT, while no significant effect observed in fruit of overexpression lines (Fig. 4B). For the three main soluble sugars (sucrose, fructose and glucose), we observed no significant change in overexpression fruit. However, in *SlVI* knockout fruit, sucrose levels were approximately sixteen times higher than in WT. Consistent with the loss of invertase activity, the levels of glucose and fructose were decreased in *SlVI* knockout fruit (Fig. 4C). Organic acids are crucial factors that contribute to the flavor and taste of fruit since they provide acidity, balance sweetness, and enhance overall flavor perception. Investigation of the main organic acids, such as citric acid and malic acid, which contribute to the unique taste profile of tomato fruit, showed no significant differences among overexpression, knockout, and WT fruit (Fig. S6). Finally, flavonoids are essential secondary metabolites in fruit and contribute to various aspects of fruit quality and potential health benefits. We measured the content of three major flavonoids and found that the knockout fruit had a nearly two-fold increase in naringenin levels compared to the wild type, but no significant difference in the levels of nicotiflorin and rutin (Fig. 4D). These data indicate that the disruption of the *SlVI* gene increases the total soluble solids and flavonoids in tomato fruit and thus contributing to improving fruit taste and nutritional value. These findings shed light on the intricate relationships between sugar metabolism, fruit taste, and metabolite profiles in tomato fruit.

**Figure 4.**
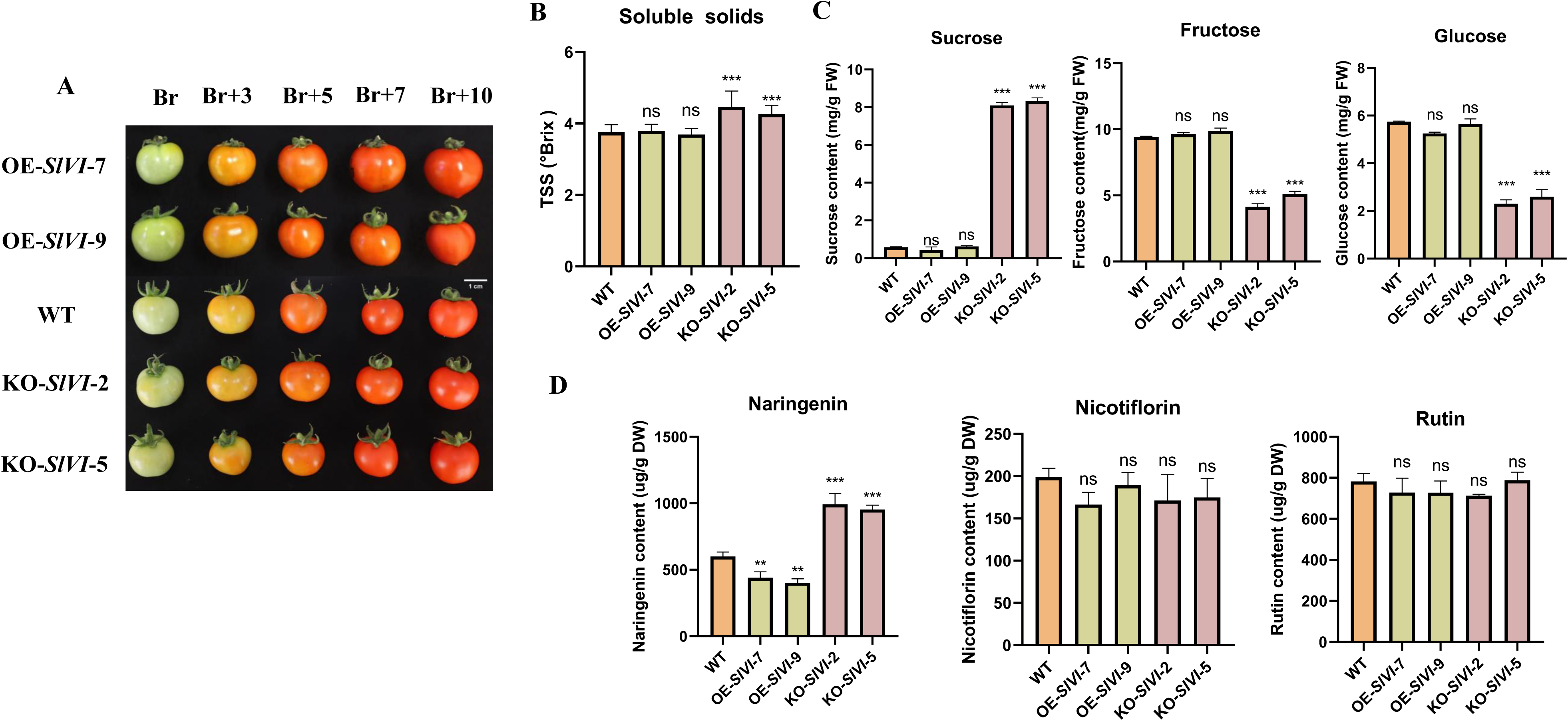
Sucrose and naringenin levels are significantly increased in postharvest *SlVI* knockout fruit. A) Phenotype of transgenic tomato fruit from breaker to ten days after breaker stages. Bar = 1 cm. B) Content of soluble solids in WT, KO-*SlVI* and OE-*SlVI* Br+5 fruit. C) Soluble sugar (sucrose, fructose, glucose) contents in WT, KO-*SlVI* and OE-*SlVI* Br+5 fruit. D) Contents of three flavonoid compounds in WT, KO-*SlVI* and OE-*SlVI* Br+5 fruit. Data are presented as means ±SD, (b, n = 20; (c-g, n=3). Asterisks indicate statistical significance using Student’s test *p < 0.05, **p < 0.01 and ***p < 0.001 and ns indicates no significant change when compared with wild type.

### Knockout of *SlVI* enhances fruit firmness by increasing the content of cell wall components

In postharvest storage, fruit firmness is a critical parameter that directly impacts the quality and shelf life of tomatoes. Maintaining optimal firmness levels helps to prevent mechanical damage, bruising, decay during transportation, handling, and storage processes. Using a texture analyzer, we measured fruit firmness at different ripening stages. The results showed that the firmness was significantly increased in *SlVI* knockout lines compared to WT throughout the ripening stages ranging from Br to 10 Br+10 (Fig. 5A). To further understand the reason for this increase in fruit firmness, we performed histological analysis on WT and *SlVI* knockout fruit. The microscopic images of the fruit’s equatorial cross-sections revealed that *SlVI* knockout fruit had a thinner pericarp compared to WT (Fig. 5B and C). However, upon counting the number of cells per five square millimeters of the pericarp, we found that the knockout fruit had a significantly higher cell density and smaller cell size compared to the WT (Fig. 5D and E). These data indicated that although the pericarp thickness of the knockout lines was significantly reduced, the cells appeared less expanded and more densely organized. The increased cell density in the knockout fruit could potentially enhance cell wall strength and overall fruit firmness.

**Figure 5.**
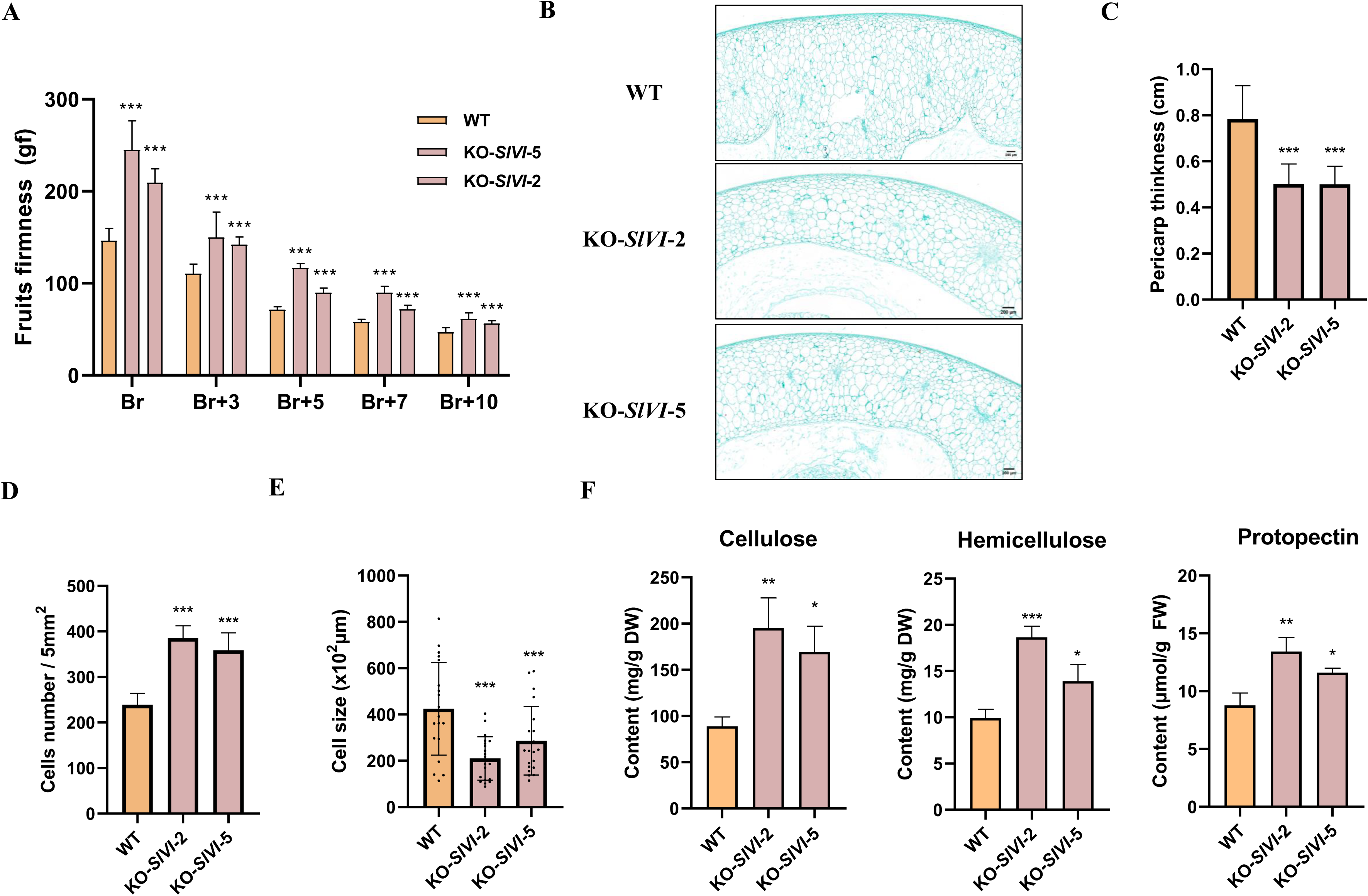
Knocking out *SlVI* improves fruit firmness and cell wall components. A) Fruit firmness in WT and KO-*SlVI* plants at different ripening stages. B) Safranin O-Fast Green staining of transverse paraffin-embedded tomato fruit sections at the Br stage in WT and KO-*SlVI*. Bars= 200 μm. C) Pericarp thickness at the Br stage in WT and KO-*SlVI*. Thickness of the pericarp in microscope images was measured using ImageJ. D) Cell number per five square millimeters in WT and KO-*SlVI* pericarps was measured using ImageJ. E) Cell size in WT and KO-*SlVI* pericarps was measured using ImageJ. F) Contents of the three main cell wall components in WT and KO-*SlVI* fruit at the Br stage. Data are presented as means ±SD, (a, e n=20. c, d, f n=3). Asterisks indicate statistical significance using Student’s test *p < 0.05, **p < 0.01 and ***p < 0.001 and ns indicates no significant change when compared with wild type.

Cellulose, hemicellulose, and protopectin are the primary components of the cell wall in most plants. They provide structural support, protection, and rigidity to the cell wall (Shi et al., 2023). To further investigate the impact of *SlVI* knocked out on cell wall composition, we quantified the content of cellulose, hemicellulose, and protopectin in the pericarp of wild type and *SlVI* knockout fruit using a spectrophotometric method. In comparison to the wild type, the pericarp of the knockout fruit displayed an approximately twofold increase in cellulose and hemicellulose content and a 1.5-fold increase in protopectin content (Fig. 5F). Increased of primary cell wall content may contributes to higher fruit firmness.

### Knockout of *SlVI* extends fruit shelf life and improves *Botrytis cinerea* resistance

We performed a shelf-life analysis to assess the postharvest quality of tomato fruit by measuring water loss (dehydration). Fruit in Br+5 stage were selected, and their weight was recorded at regular intervals during a storage period of 50 days at room temperature. Interestingly, we found that *SlVI* knockout fruit exhibited less water loss symptoms, characterized by visible wrinkling and collapsing, indicating an improve in shelf life compared with WT fruit (Fig. 6A). The weight loss data also revealed that WT fruit exhibited a higher rate of weight loss starting from 30 days post-harvest, indicating extended shelf life of *SlVI* knockout fruit compared to WT (Fig. 6B). To further illustrate the water loss phenotype, we stained sections of fruit to visualize the cuticle thickness. As shown in Fig. 6C, D, *SlVI* knockout fruit have thicker cuticles compared to wild type. Taken together, our findings suggest that knockout of *SlVI* positively influences fruit firmness by increasing the pericarp cell density and cell wall polymer content. In addition, knockout significantly extended the shelf life of the fruit likely by increasing the thickness of the cuticle.

**Figure 6.**
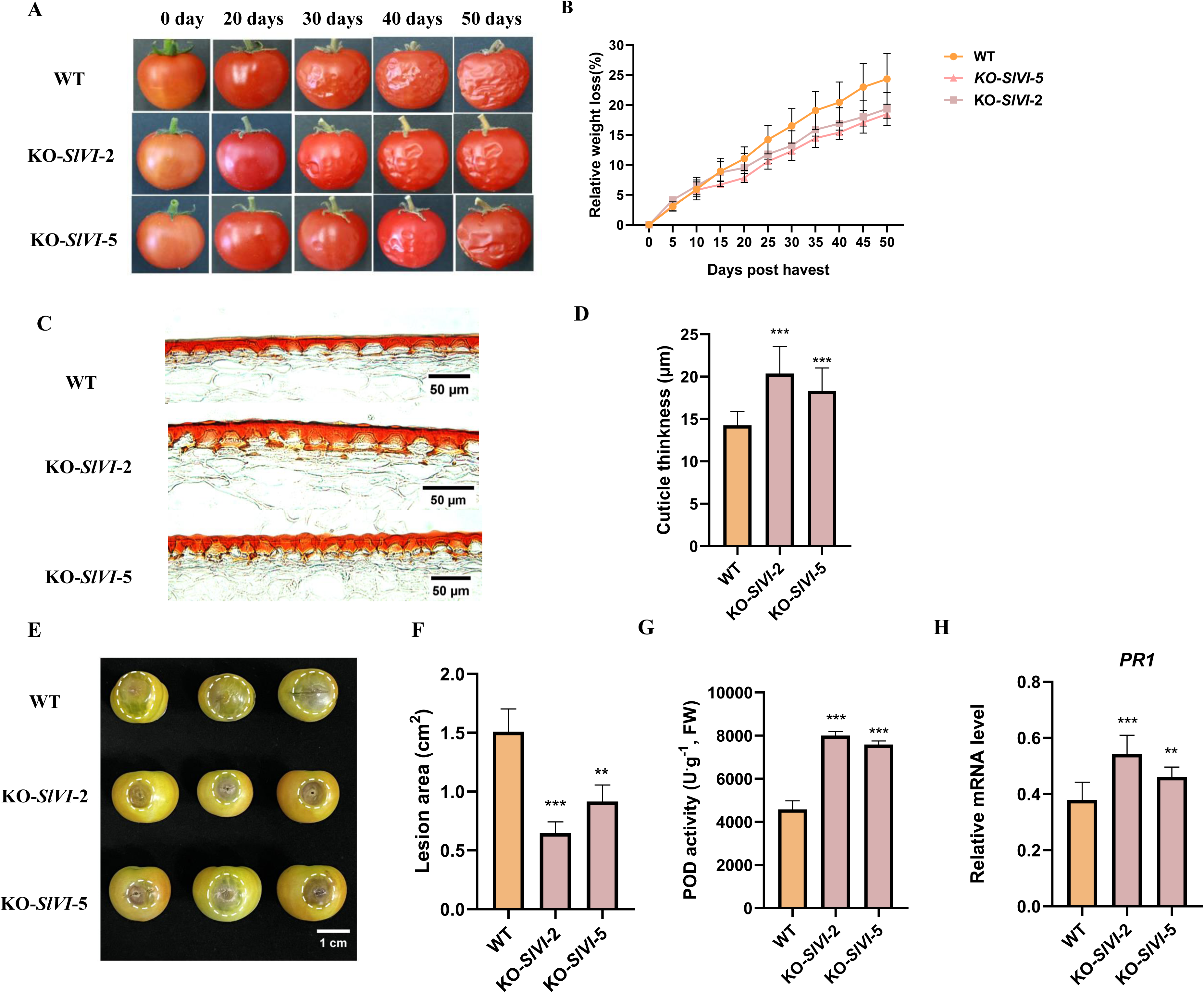
Knockout of *SlVI* extends fruit shelf life and improves fruit *Botrytis cinerea* resistance. A) Phenotype of WT and KO-*SlVI* fruit during storage. B) Relative weight loss (weight/fresh weight) in WT and KO-*SlVI* fruit during storage. C) Micrographs of fruit epicarp transverse sections at Br stage in WT and KO-*SlVI* lines to visualize the cuticle. Bars = 50 μm. D) Fruit cuticle thickness in WT and KO-*SlVI* lines. Cuticle thickness was measured by ImageJ. E) Symptoms of WT and KO-*SlVI* fruit 48 hours post inoculation (hpi) with *B. cinerea.* White circles indicate the lesion margins. F) Lesion in WT and KO-*SlVI* fruit after infection. Lesion areas was measured by ImageJ. G) Antioxidant peroxidase activity in WT and KO-*SlVI* tomato fruit after inoculation with *B. cinerea*. H) Relative expression levels of the disease resistance related *PR1* gene in WT and KO-*SlVI* fruit after infection.

Improving the postharvest storability of fruits and resistance to both biotic and abiotic stress is an important strategy to reduce food waste and economic losses in the fresh fruit industry. To evaluate the resistance of *SlVI* knockout fruit to pathogen infections, we conducted a *B. cinerea* inoculation experiment. At the Br stage, the fruit surfaces were inoculated with *B. cinerea*. After 48 hours, we observed that WT fruit produced larger lesion areas compared with KO-*SlVI*-2 and KO-*SlVI*-5, indicating increased resistance to *B. cinerea* in the *SlVI* knockout fruit (Fig. 6E and F). Furthermore, we analyzed peroxidase (POD) activity and the expression of *SlPR1*, an established pathogen response gene, in fruit after *B. cinerea* infection. The results showed an increase in both POD activity and expression of *SlPR1* in *SlVI* knockout fruit, while these remained relatively low in WT fruit (Fig. 6G, H). These findings suggest that knocking out *SlVI* gene in tomato enhances fruit resistance against *B. cinerea* infection, possibly through the activation of defense-related genes. These findings demonstrate that the alteration of sugar metabolism by *SlVI* knocked out can confer improved resistance to *B. cinerea*.

### Transcriptome analysis revealed changes of sugar and postharvest quality-related genes in *SlVI* knockout fruit

To gain a deeper insight into how *SlVI* knockout affects tomato fruit sugar metabolism and fruit postharvest quality at molecular levels, we conducted global gene expression profiling using RNA-seq analysis of KO-*SlVI*-2 and WT fruit at Br+5 stage. In total, we detected 3280 differentially expressed genes (DEGs), of which 1655 were upregulated, and 1625 were downregulated in KO-*SlVI*-2 compared to WT. (Fig. 7A and Data S1). Kyoto Encyclopedia of Genes and Genomes (KEGG) annotation of these DEGs indicated that multiple metabolic pathways are significantly affected in *SlVI* knockout fruit. Genes involved in photosynthesis, phenylpropanoid and flavonoid biosynthesis, starch and sucrose metabolism, as well as cutin and wax biosynthesis were enriched in the DEGs (Fig. 7B). A gene ontology (GO) analysis also indicated that knockout of *SlVI* affected multiple biological process, including the carbohydrate metabolic process, response to oxidative stress, cell growth and cell wall macromolecule catabolic process. In addition, some molecular functions, such as iron ion binding, DNA-binding transcription factor activity, and transmembrane transporter activity are also affected (Fig. S7).

**Figure 7.**
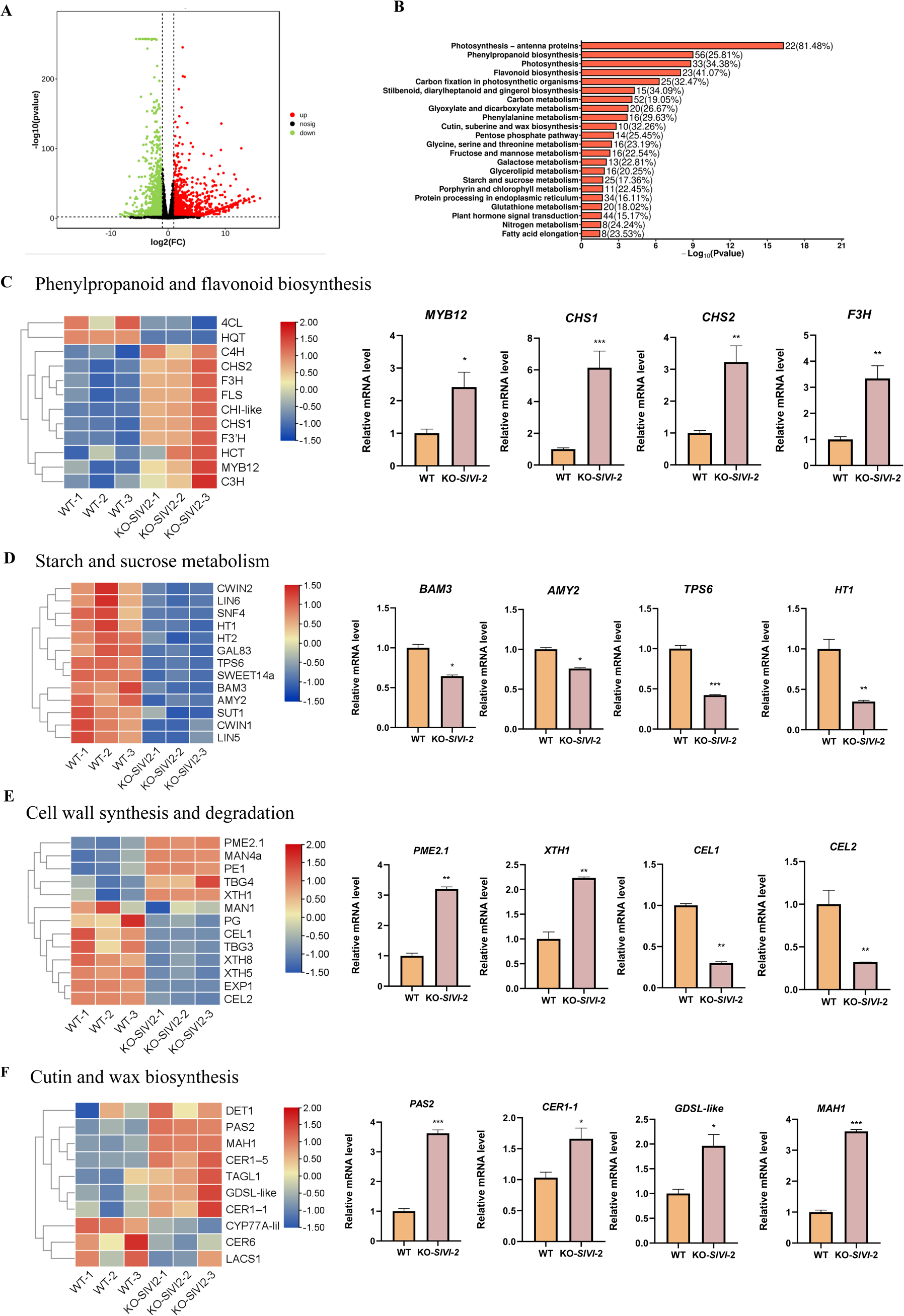
RNA-seq profiling of WT and KO-*SlVI* fruit. A) Volcano plot displaying the differential gene expression profiles in WT and KO-*SlVI* samples. B) Kyoto Encyclopedia of Genes and Genomes (KEGG) annotation of DEGs between WT and KO-*SlVI*-2. C) Heatmap and RT-qPCR analysis of phenylpropanoid and flavonoid biosynthesis gene expression patterns. D) Heatmap and RT-qPCR analysis of starch and sucrose metabolism related gene expression patterns. E) Heatmap and RT-qPCR analysis of cell wall synthesis and degradation related gene expression patterns. F) Heatmap and RT-qPCR analysis of cutin and wax biosynthesis related gene expression patterns. Data are presented as means ± SD, (n=3). Asterisks indicate statistical significance using Student’s test *p < 0.05, **p < 0.01 and ***p < 0.001 and ns indicates no significant change when compared with wild type.

We first verified the impact on the expression of genes related to the metabolism of flavonoid, starch and sucrose. As shown in heat map, the trans-cinnamate 4-monooxygenase (*4CH*) gene, which is involved in the phenylpropanoid metabolic pathway, shows higher expression levels in fruit of *SlVI* knockout lines compared to WT (Fig. 7C). Transcription factor *SlMYB12*, which plays a key role in regulating the downstream flavonoid metabolic pathway, also exhibits higher expression level in the knockout fruit. Moreover, the higher expression levels of chalcone synthase genes (*CHS1* and *CHS2*) in the knockout fruit aligns with the increase in naringenin chalcone content. Furthermore, genes (*F3H* and *FLS*) related to the downstream steps in the flavonoid biosynthesis pathway are also highly expressed in *SlVI* knockout fruit (Fig. 7C). This further supports the notion that disruption of the *SlVI* gene resulted in an increase in the production of flavonoid compounds (Fig. 4D). In addition, we also focused on the expression of genes related to starch degradation and sugar transport aimed to characterize the dynamic energy transport and transformation processes in fruit. The heatmap analysis indicated a decrease in the expression of two genes, *BAM3* (Beta-amylase 3) and *AMY2* (Alpha-amylase 2), which play pivotal roles in the hydrolysis of starch into maltose and shorter oligosaccharides, respectively (Fig. 7D). This decline could potentially be attributed to the substantial accumulation of sucrose observed in *SlVI* knockout fruit, leading to negative feedback mechanism in energy conversion pathways. Furthermore, a significant decrease in the expression of *TPS6* (encoding a trehalose-6-P synthase-like protein) and *HT1* (encoding a hexose transporter) was observed in *SlVI* knockout fruit (Fig. 7D).

To investigate the changes in fruit texture and firmness at molecular levels, we examined the expression levels of cell wall synthesis and degradation related genes. The *SlVI* knockout fruit exhibited increased expression levels of xyloglucan endotransglucosylases/hydrolases (*XTHs*) and pectin methylesterases (*PMEs*) (Fig. 7E), suggesting an enhanced cell wall synthesis in *SlVI* knockout fruit. In contrast, the expression levels of poly-galacturonases (*PGs*) and cellulases (*CEL1*, and *CEL2*) involved in cell wall degradation were decreased in knockout fruit (Fig. 7E). This data is consistent with the extended shelf life and higher firmness observed in the knockout fruit (Fig. 5A, 6A and B). In response to the remarkable reduction in water loss rate observed in knockout fruit during postharvest storage, we checked the expression of genes associated with the fruit cuticle wax biosynthesis. Notably, the expression levels of *SlCER1-1*, a key gene involved in the biosynthesis of the precursor for cuticular wax components, were higher in *SlVI* knockout fruit compared to WT (Wu et al., 2022). Additionally, *PAS2*, known to function in lipid metabolism or cuticle remodeling and, contributing to the formation and maintenance of the cuticular in tomato fruit, also exhibited increased expression in the knockout fruit (Barraj Barraj et al., 2021) (Fig. 7F). These results indicate that sugar transport in fruit is a highly dynamic and balanced process, where changes in one gene can trigger a cascade of alterations in other genes involved in the different metabolic pathways.

### Transcriptional regulation of *SlVI* by key ripening regulators

To investigate whether key ripening-related transcription factors can regulate the expression of the *SlVI* gene, we conducted a dual luciferase reporter assay. The results showed that transcription factors RIN, FUL1, and NOR can significantly activate the expression of *SlVI*, with activation levels approximately 2 to 5 times higher than the control. Furthermore, when RIN and FUL1 were co-expressed with the *SlVI* promoter, the activation effect increased to 30 times, suggesting a synergistic effect rather than a mere additive effect. However, we observed that RIN and NOR, as well as NOR and FUL1, do not exhibit synergistic effects (Fig. 8A and B). Yeast two-hybrid experiments further revealed that RIN and FUL1 can interact with each other, but not for NOR and FUL1 (Fig. 8C). To further support the conclusion that the specific expression of the *SlVI* gene during fruit ripening is regulated by ripening-related transcription factors, we examined the expression of the *SlVI* in rin, nor mutants, and co-silenced *FUL1/2* fruit. The results showed that the relative expression of *SlVI* was significantly decreased in these mutants compared to WT (Fig. 8D), supporting a regulatory role of these key ripening regulators on sugar metabolism by regulation of *SlVI*.

**Figure 8.**
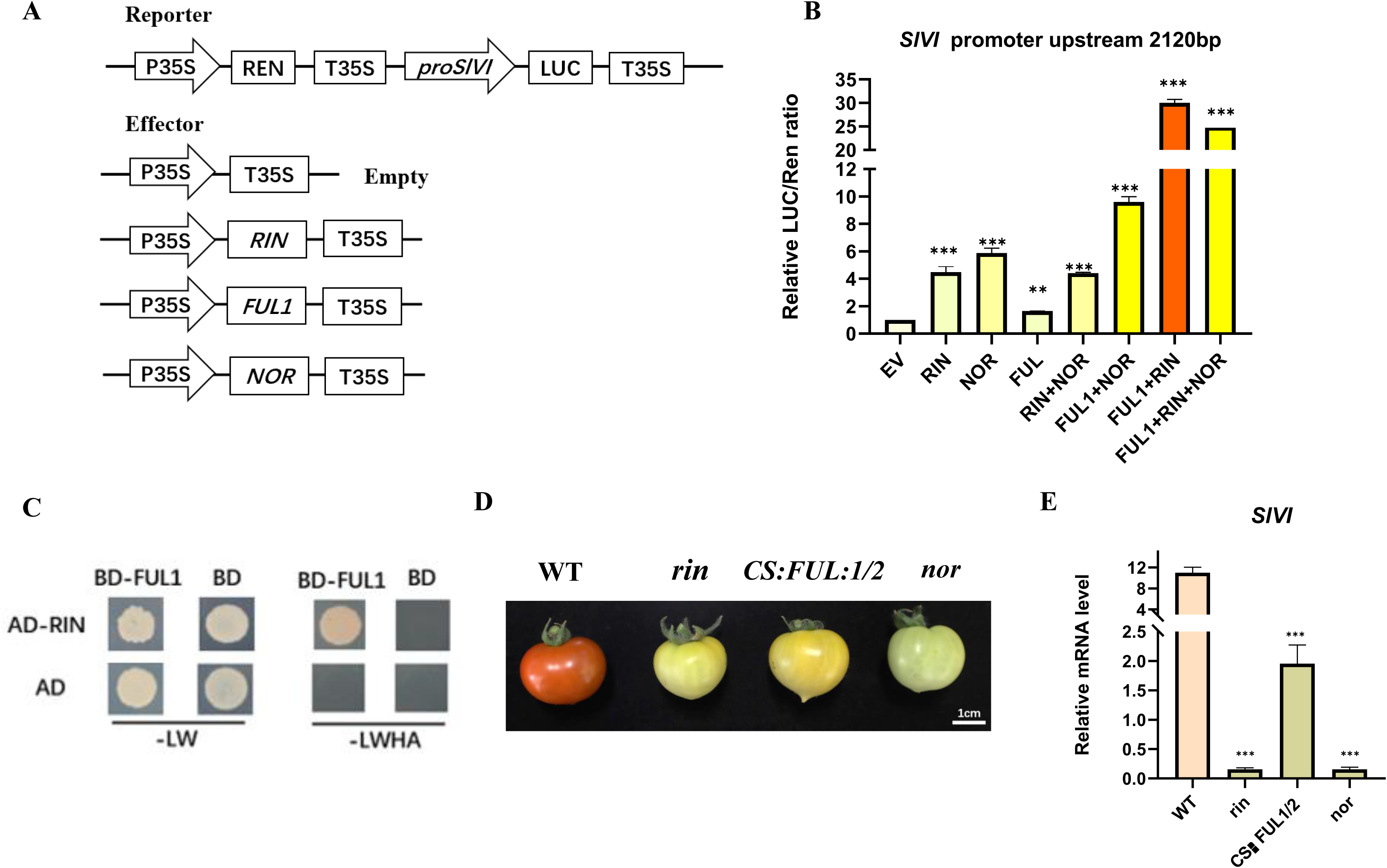
Transcriptional regulation of SlVI by key ripening regulators. A) A schematic illustration of the LUC reporter and effector constructs used in transient expression assays. B) The activation of *SlVI* by ripening related transcription factors. LUC, firefly luciferase; REN, Renilla luciferase. C) Analysis of the interaction between RIN and FUL1 in yeast system. D) Phenotype of *rin*, *nor* mutant and co-silence *FUL1/2* Br+5 fruit. E) Relative expression level of *SlVI* genes in rin, nor mutant and co-slience *FUL1/2* fruit. Data are presented as means ± SD, (n=3). Asterisks indicate statistical significance using Student’s test *p < 0.05, **p < 0.01 and ***p < 0.001 and ns indicates no significant change when compared with wild type.

## Discussion

Sucrose not only serves as a vital carbon and energy source for plant growth and development but also participates in numerous biochemical reactions through a series of metabolic processes (Wang et al., 2001; Roitsch and González, 2004). Additionally, sugars such as sucrose, glucose and fructose also act as signaling molecules that can more directly regulate various physiological and developmental processes in response to changing environmental conditions (Koch, 2004; Lastdrager et al., 2014). In our study, we focused on the vacuolar invertase SlVI, which hydrolyses sucrose into glucose and fructose is induced during tomato fruit ripening. We observed a remarkable increase in sucrose and decrease in glucose and fructose content in *SlVI* knockout fruit without significant changes in the fruit ripening process (Fig. 4A, 4C, S5). For flavonoid metabolism pathways, *SlVI* knockout fruit exhibiting a nearly two-fold increase in naringenin content compared to the wild type (Fig. 4D). From the transcriptome data, it is evident that many genes involved in the phenylpropanoid and flavonoid metabolic pathways are significantly up-regulated in knockout fruit (Fig. 7C). These findings align with previous studies demonstrating that sucrose promotes the accumulation of flavonoids in *Arabidopsis*, which may be mediated by DELLA protein stabilization (Solfanelli et al., 2006; Li et al., 2013). Plant cell wall metabolism is a multifaceted process, intricately regulated by various genes with distinct functions, even within the same gene family (Shi et al., 2023). Texture analysis revealed a significant increase in fruit firmness in knockout fruit and higher relative content of cell wall components in pericarp tissue compared to WT (Fig. 5A, F). According to previous studies, overexpressing *SlXTH1* or knockout of *SlXTH5* in tomato can increase fruit firmness, while silencing of *SlPME2.1* in tomato fruit resulted in an increased rate of softening during ripening, which is consistent with our transcriptome data related to cell wall synthesis and degradation (Fig. 7E) (Miedes et al., 2010; Wang et al., 2023). Moreover, simultaneous inhibition of *SlPG* and *SlEXP1*, or inhibition of *SlPL*, has been demonstrated to reduce cell wall disassembly and elevate levels of cellulose and hemicellulose. This impedes fruit softening and enhances resistance to *B. cinerea* (Cantu et al., 2008; Yang et al., 2017). In our *SlVI* knockout fruit the substantial downregulation of these genes, as depicted in the heatmap, may closely correlate with the increased content of cell wall components and the improved resistance to *B. cinerea* (Fig. 6E,7E). Besides, increased content of flavonoids in tomato fruit also has a positive effect on resistance to *B. cinerea* infection (Zhang et al., 2013). This suggests a synergistic interplay between flavonoids and cell wall composition in contributing to the enhanced resistance of *SlVI* knockout fruit against *B. cinerea*. Histological analysis demonstrated that the knockout fruit exhibited smaller but densely arranged cells (Fig. 5B, 5D, 5E). While there are relatively few reports directly linking endogenous sugar content to cell development and expansion, it is well-established that sugar can exert regulatory effects on these processes by modulating plant hormone levels (León and Sheen, 2003; Sairanen et al., 2012). From our transcriptome data and RT-qPCR validation, it can be concluded that key genes involved in multiple hormones signaling pathways are altered in knockout fruit (Fig. S8). However, the precise molecular mechanisms underlying these observations require further investigation.

Consistent with previous studies highlighting the role of sucrose signaling in regulating the transition from the juvenile to the adult phase in *Arabidopsis*, the *SlVI* overexpression lines, characterized by increased hexose levels in leaves, exhibited an early flowering phenotype (Meng et al., 2021). Additionally, we investigated the expression level of *SlSBP3*, a gene homologous to an *Arabidopsis* know positive regulator of flowering (Silva et al., 2018). As expected, the expression level of *SlSBP3* also increased in the *SlVI* overexpression lines (Fig. 3B, G, H). These findings suggest that altering sugar metabolism can accelerate the vegetative growth stage and promote early flowering, possibly through altered hexokinase levels and signaling.

In conclusion, our study highlights the importance of sucrose metabolism in tomato fruit development, flavor, texture, and postharvest quality. Disruption of the *SlVI* gene increased sucrose and naringenin content and improved fruit firmness by increasing endocarp cell wall polymer content, and coincided with enhanced resistance to *B. cinerea* infection. These findings expand our understanding of the molecular mechanisms underlying fruit development and ripening processes regulated by sucrose metabolism. Further research is warranted to unravel the intricate regulatory networks and molecular interactions involved in sugar and energy sensing and signaling and the modulation of fruit quality traits. The findings of this study provide valuable insights for the development of strategies aimed at improving fruit quality and postharvest storage of tomatoes.

## Materials and methods

### Plant material and growth conditions

Tomato plants (*Solanum lycopersicum* cv Micro Tom) were cultivated in a controlled greenhouse environment following standard culture conditions (16/8 h of light/dark cycle at 24°C in 60% humidity and under 250 μmol/m^-2^/s^-1^ intense luminosity) and watered daily. For sample collection, various tomato tissues and different developmental stages of tomato fruit (without seeds) were carefully harvested. The samples were immediately frozen in liquid nitrogen to ensure rapid preservation of their metabolic compounds and RNA content. Subsequently, the frozen samples were stored at −80 °C in a freezer until further processing.

### Plasmid construction and plant transformation

The full coding sequence (CDS) of the *SlVI* gene (*Solyc03g083910*) was integrated into the binary plant expression vector pBI121. The pBI121 vector was chosen for its constitutive CaMV35S promoter, which allows for strong and continuous expression of the *SlVI* gene. Additionally, the vector contained a kanamycin resistance gene, enabling the selection of transgenic plants through antibiotic resistance. To facilitate genome editing and generate knockout, the CRISPR/Cas9 vector, named BGFastCas9-plant, was obtained from BioGround Corp company and modified accordingly. Dr. Xian Zhiqiang from BioGround Corp company in Chongqing, China kindly provided the modified vector for this study, granting permission for its use (Vazquez-Vilar et al., 2016). The CRISPR-P 2.0 online tool (http://cbi.hzau.edu.cn/crispr/) was utilized to design a specific target site for the *SlVI* gene. Primers required for the construction of the CRISPR/Cas9 system are listed in Table S1. The constructed *SlVI* overexpression and CRISPR/Cas9 vectors were introduced into Agrobacterium tumefaciens strain GV3101. The transformation of tomato plants was performed using Agrobacterium-mediated methods (Ying et al., 2020).

### RNA extraction and RT-qPCR analysis

Total RNA was extracted from various tomato tissues using a plant RNA extraction kit (BIOFIT, Chengdu, China). First-strand cDNA was reverse transcribed from 1 μg of total RNA using the PrimeScriptTM RT reagent kit (AK4201; Takara Bio, Kusatsu, Japan) according to the manufacturer’s instructions. For quantitative real-time PCR (RT-qPCR), the BG0014 SYBR Prime qPCR Set (Fast HS; BioGround Corp company, Chongqing, China) was employed. The RT-qPCR experiments were performed on a Bio-Rad CFX96 Real-Time PCR System. The RT-PCR program and data analysis procedures followed the established method as described in previous studies by You et al. The primer sequences used for RT-qPCR in this study are provided in Supplemental Table S1 and using Actin (*Solyc11g005330*) as the reference gene. All tissues were obtained from tomato plants grown under the same growth conditions, and each sample included three technical replicates.

### Chemical compound content analysis

To extract starch from tomato fruit powder, a starch detection kit from Solarbio, Beijing, China, was used following the manufacturer’s provided instructions. For the preparation of sugar standard solutions, fructose, glucose, and sucrose were individually dissolved in a mixture of 50% acetonitrile (ACN)/H2O at various concentrations ranging from 0.5 to 1 mg/mL. Similarly, citric acid and malic acid standards were dissolved in a 50% methanol/H2O solution at the same concentrations. Tomato fruit powder was first dissolved in the same solution with standards at a ratio of 0.1 g/mL and diluted 100 times and filtered using a 0.45 µm PVDF syringe filter for analysis. All analyses were performed using the SCIEX Triple Quad 5500 LC-MS/MS System. For sugar and organic acid analysis, ACQUITY UPLC BEH Amide Columns (1.7 µm, 2.1 x 100 mm, made in Ireland) were used to separate and quantify the compounds. Flavonoid metabolites were quantified following the method described by (Yuan et al., 2022). All standards used in these analyses were purchased from Shanghai Yuanye Bio-Technology (Shanghai, China).

### Fruit firmness and shelf life tests

To measure fruit firmness, a TA.XTC-18 texture analyzer (Bosin Tech, Shanghai, China) was utilized with a P/2 columnar probe of 2 mm diameter. More than twenty fruit at the Br+5 stage were selected for the firmness measurements. The test parameters and calculation method were carried out as previously described in the study by You et al. For shelf life testing, fruit at the Br+5 stage were harvested and placed in a plant climate chamber set at room temperature conditions (25 ℃, 55–60% relative humidity). The physiological loss of water (PLW) was calculated by measuring the weight loss per fruit every 5 days of storage. The appearance and quality of the fruit were visually assessed and photographed every 10 days throughout the storage period. At least ten fruit were collected from each transgenic line for analysis and comparison.

### Cytological assessment and main cell wall components content determination

Tomato Br stage fruit was harvested for Safranin O-Fast Green staining (G1101, Wuhan Servicebio Technology Co., Ltd, China) and the method is consistent with the previous description (You et al., 2024). For the three main cell wall components content, cellulase, hemicellulose and protopectin content assay kit were purchased from Solarbio (Beijing Solarbio Science & Technology Co.,Ltd) and using the spectrophotometer according to the manufacturer’s instructions.

### *Botrytis cinerea* resistance analysis

The pathogen *Botrytis cinerea* was collected and growth on PDA medium at 20 °C. Tomato fruit at Br stage was inoculated with mycelial plugs and storage at room temperature with relative high humidity for phenotype observation and further experimentation (Pei et al., 2019). Infection symptoms were photographed and calculated at 48 h after inoculated using Image J.

### Transcriptome profiling

In our study, we performed RNA sequencing (RNA-seq) to analyze the transcriptome profiles of tomato fruit at the Br+5 stage for both the wild-type and KO-*SlVI*-2 lines. RNA-Seq library construction and high-throughput sequencing were conducted by Biomarker Technologies (Beijing, China). The library preparations were sequenced on an Illumina Novaseq 6000 platform, and 150 bp paired-end reads were generated. The library quality was then assessed using an Agilent Bioanalyzer 2100 system and sequenced on an Illumina Hiseq X-Ten platform to obtain raw reads. Transcriptome analysis was carried out as previously described (Yuan et al., 2022). DEGs were found using a significance threshold of a log 2 fold change of ±1. KOBAS software was used to test the statistical enrichment of DEGs in KEGG pathways. Pathways with the P < 0.05 were significantly enriched. Gene Ontology (GO) analysis is a method used gene annotations from a database Gene Ontology Consortium (http://geneontology.org/).

### Yeast two-hybrid assay

The full-length coding sequences of *RIN*, *NOR*, and *FUL1* were cloned into the pGADT7 and pGBKT7 vectors, respectively. These constructs were then transformed into the AH109 yeast strain. Yeast transformants were cultured on Synthetic dropout (SD) medium lacking Leu and Trp (SD-Leu-Trp) for 2–4 days to select for positive transformants.To confirm the interaction between the two proteins, selection medium plates (-Leu, -Trp, -His, -Ade) were utilized. Growth of yeast colonies on these plates indicated an interaction between the proteins.

### Dual-luciferase reporter assay

The promoter sequence of the *SlVI* (*Solyc03g083910*) gene was obtained from the tomato genome website (https://solgenomics.net/). This promoter sequence was then constructed into a dual luciferase reporter vector containing a firefly luciferase reporter gene (Luc) and a CaMV35S promoter-driven Renilla luciferase reporter gene (Ren), with Ren serving as an internal reference reporter gene. The CDS sequences of the maturation-related transcription factors *RIN*, *NOR*, and *FUL1* were cloned into a plant expression vector as effectors.Using transient transformation of tobacco leaves and the Dual-Luciferase reporter assay system (Promega, Madison, WI, USA), we tested the ratio of LUC to REN to determine the activation of the *SlVI* promoter by these transcription factors.

### Statistics

All data are expressed as mean ± SD from 3 or more independent experiments and subjected to Student’s t test for pairwise comparison or ANOVA for multivariate analysis.

## Author contributions

M.L., Y.Wu., Y.Wang. and N.H planned and designed the research; Y.W., H.C., M.W., Y.Z., C.Y. and Q.Y. performed experiments and analyzed data. M.L. and Y. Wu. wrote the manuscript, M.B., F.R. and B.V.d.P. helped improve the manuscript.

## Supplemental data

Additional supporting information may be found online in the Supporting Information section at the end of the article.

**Figure S1** Tomato fruit at different developmental stages were stained with iodine solution.

**Figure S2** Phloem unloading of sucrose (S) in sinks may occur either apoplasmically into cell wall matrix or symplasmically through plasmodesmata (PD) in recipient cells.

**Figure S3** Heat map of sucrose transport and metabolism genes expression levels from TEA *S. lycopersicum* M82 and Micro Tom RNA-seq data.

**Figure S4** Phylogenetic analysis of SlVI and related proteins from *Arabidopsis thaliana* and tomato.

**Figure S5** Time from anthesis to breaker stage in WT, KO-*SlVI* and OE-*SlVI* lines.

**Figure S6** Malic acid and citric acid contents in WT, KO-*SlVI* and OE-*SlVI* Br+5 fruit.

**Figure S7** A gene ontology (GO) analysis of DEGs between WT and KO-*SlVI*-2.

**Figure S8** RT-qPCR in KO-*SlVI*-2 and wild type fruit at Br+5 stages of genes involved in plant hormones pathway.

**Table S1** Primers used in this study.

**Data S1** All sample RNA-seq TPM, DEGs enrichment in KEGG pathways and GO analysis.

## Data availability

The data that support the findings of this study are available from the corresponding author, Mingchun Liu, upon reasonable request.

